# Annihilation of Methicillin-resistant *Staphylococcus aureus* via Photobleaching of Staphyloxanthin

**DOI:** 10.1101/227603

**Authors:** Pu-Ting Dong, Haroon Mohammad, Xiaoyu Wang, Jie Hui, Junjie Li, Lijia Liang, Mohamed N. Seleem, Ji-Xin Cheng

**Affiliations:** Department of Chemistry, Boston University, Boston, MA 02215, USA; Department of Comparative Pathobiology, College of Veterinary Medicine, Purdue University, West Lafayette, IN 47907, USA; Molecular Biology, Cell Biology & Biochemistry, Boston University, Boston, MA 02215, USA; Department of Physics and Astronomy, Purdue University, West Lafayette, IN 47907, USA; Department of Electrical and Computer Engineering, Boston University, Boston, MA 02215, USA; State Key Laboratory of Supramolecular Structure and Materials, Institute of Theoretical Chemistry, Jilin University, Changchun 130012, China; Department of Biomedical Engineering, Boston University, Boston, MA 02215, USA; Photonics Center, Boson University, Boston, MA 02215, USA

**Author notes:** To whom correspondence should be addressed: Ji-Xin Cheng and Mohamed N. Seleem.

## Abstract

Given that the dearth of new antibiotic development loads an existential burden on successful infectious disease therapy^1^, health organizations are calling for alternative approaches to combat methicillin-resistant *Staphylococcus aureus* (MRSA) infections. Here, we report a drug-free photonic approach to eliminate MRSA through photobleaching of staphyloxanthin, an indispensable membrane-bound antioxidant of *S. aureus*^2-5^. The photobleaching process, uncovered through a transient absorption imaging study and quantitated by absorption spectroscopy and mass spectrometry, decomposes staphyloxanthin and sensitizes MRSA to reactive oxygen species attack. Consequently, staphyloxanthin bleaching by low-level blue light eradicates MRSA synergistically with external or internal reactive oxygen species. The effectiveness of this synergistic therapy is validated in MRSA culture, MRSA-infected macrophage cells, *S. aureus* biofilms, and a mouse wound infection model. Collectively, these findings highlight broad applications of staphyloxanthin photobleaching for treatment of MRSA infections.

---

*Staphylococcus aureus* causes a variety of diseases ranging from skin and soft tissue infections to life-threatening septicemia^6-9^. Moreover, *S. aureus* has acquired resistance to multiple antibiotic classes that were once effective^10^. A classic example is the emergence of clinical isolates of methicillin-resistant *Staphylococcus aureus* (MRSA) strains in the 1960s that exhibited resistance to *β*-lactam antibiotics^11-13^. More recently, strains of MRSA have exhibited reduced susceptibility to newer antibiotics such as daptomycin and antibiotics deemed agents of last resort such as vancomycin and linezolid^14,15^. Besides the acquired resistance through mutational inactivation, MRSA develops other strategies, e.g. residing inside host immune cells or forming biofilms, to evade the effect of antibiotics. Those strategies pose an appalling challenge to the successful therapy for MRSA infections.

Initially we attempted to differentiate MRSA from methicillin-susceptible *S. aureus* by transient absorption imaging (see Methods and Supplementary Fig. 1) of their intrinsic chromophores. Intriguingly, once the cultured *S. aureus* was placed under the microscope, the strong signal measured at zero delay between the 520-nm pump and 780-nm probe pulses quickly attenuated over second time scale (Fig. 1a and Supplementary Video 1). We hypothesized that a specific chromophore in *S. aureus* is prone to photobleaching under the abovementioned setting. To verify the photobleaching phenomenon, we fitted the time-course curve with a photobleaching model^16^ (Fig. 1b):

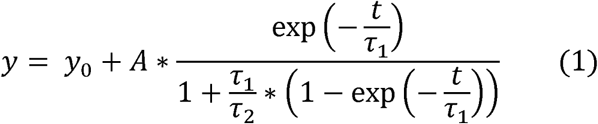

where *t* is the duration of light irradiation, y is the signal intensity, *y_0_* and A are constants, *τ*_1_ and *τ*_2_ are the time constants for the first-and second-order bleaching, respectively. The firstorder bleaching occurs at low concentration of chromophores (singlet oxygen involved, *τ*_2_ = ∞). The second-order bleaching takes place when quenching within high-concentration surrounding chromophores dominates (*τ*_1_ = ∞, Supplementary Fig. 2). Derivation of equation (1) is detailed in Methods. Strikingly, this photobleaching model fitted well the raw time-course curve (*τ*_1_ = ∞, *τ*_2_ = 0.15 ± 0.02 s, R^2^ = 0.99). Moreover, oxygen depletion (Na_2_S_2_O_4_: oxygen scavenger) showed negligible effect on the bleaching speed (*τ*_2_ = 0.14 ± 0.01 s, Supplementary Fig. 3a). The same phenomenon was observed in methicillin-susceptible *S. aureus* (Supplementary Fig. 3b). Collectively, these data support a second-order photobleaching process.

**Fig. 1.**
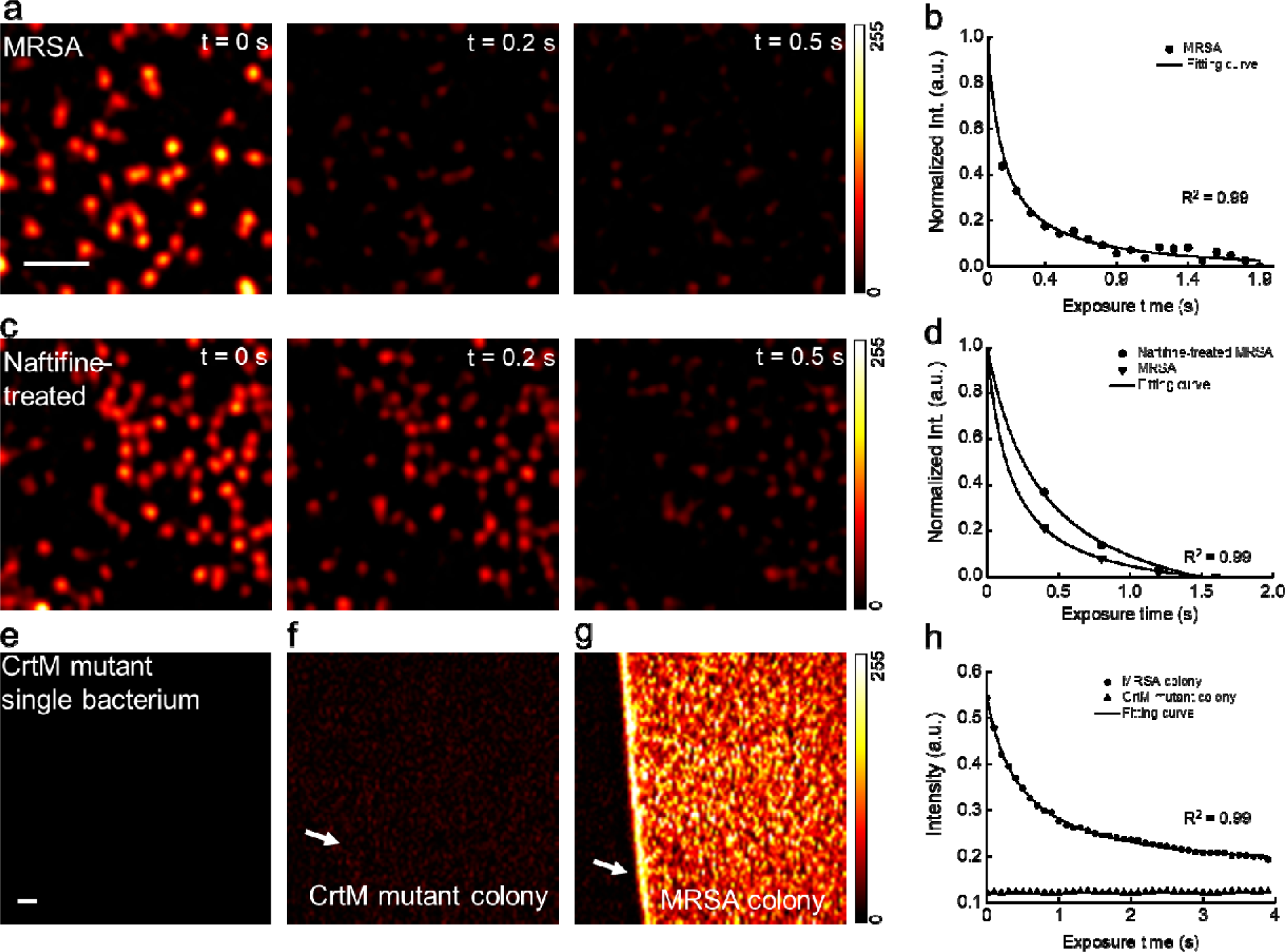
Photobleaching of staphyloxanthin in MRSA uncovered by transient absorption microscopy. (a) Time-lapse images of MRSA. Scale bar, 5 m, applies to (a,c). (b) Representative normalized time-lapse signal from MRSA. (c) Time-lapse images of naftifine-treated MRSA. (d) Representative normalized time-lapse signals from MRSA and naftifine-treated MRSA. (e-g) Images of CrtM mutant, CrtM mutant colony, MRSA colony at t = 0 s, respectively. Scale bar, 20 m, applies to (e-g). (h) Representative raw time-lapse signals from MRSA colony and CrtM mutant colony. White arrows indicate the interface between air and sample. Time-lapse signals were fitted by equation (1). The images are processed from the raw data with dynamic range of 0-255 through ImageJ.

Next, we aimed to deduce the specific chromophore inside *S. aureus* that accounts for the observed photobleaching phenomenon. It is known that carotenoids are photosensitive due to the conjugated C=C bonds^17,18^. Therefore, we hypothesized that staphyloxanthin (STX), a carotenoid pigment residing in the cell membrane of *S. aureus,* underwent photobleaching in our transient absorption study. To test this hypothesis, we treated MRSA with naftifine, a FDA-approved antifungal drug that blocks the synthesis of STX^3^. The treated MRSA exhibited lower signal intensity (Fig. 1c) and slower photobleaching speed (Fig. 1d). Specifically, *τ*_2_ of naftifine-treated MRSA (0.39 ± 0.07 s) is 2.5 times of that of MRSA (0.15 ± 0.02 s), in consistence with second-order photobleaching. Furthermore, no transient absorption signal was observed in a *S. aureus* stain containing a mutation in dehydrosqualene synthase (CrtM) (Fig. 1e) that is responsible for STX biosynthesis^19^. To avoid the systematic error aroused by single bacterium measurement, we repeated the same analysis using bacterial colonies. It turned out that CrtM-mutant colony (Fig. 1f,h) only exhibited background induced by cross-phase modulation^20^, whereas the MRSA colony showed a sharp contrast against the background (Fig. 1g) and a fast photobleaching decay (Fig. 1h). Taken together, these data confirm that STX in *S. aureus* accounts for the observed photobleaching.

In the transient absorption study, when changing 520-nm pump irradiance while fixing 780-nm probe intensity, both signal intensity and *τ*_2_ changed drastically (Supplementary Fig. 4a,c), whereas the alteration of probe irradiance only affected the transient absorption signal intensity but not *τ*_2_ (Supplementary Fig. 4b,d). These findings collectively imply that photobleaching efficacy is highly dependent on the excitation wavelength (Supplementary Fig. 4e), which is consistent with the fact that photobleaching is grounded on the absorption of chromophore^21^.

To find the optimal wavelength for bleaching STX, we measured the absorption spectrum of crude STX extract from *S. aureus*. The extract shows strong absorption in the window from 400 nm to 500 nm (Fig. 2a). Based on this result, we built a portable device composed of a blue light-emitting diode (LED) with central emission wavelength around 460 nm for wide-field bleaching of STX (Fig. 2a and Supplementary Fig. 5). We exposed the crude STX extract to blue light (90 mW) for different time intervals. Remarkably, the distinctive golden color of STX disappeared within 30-min exposure, whereas the control group under ambient light remained unchanged (Fig. 2b). Its absorption peak between 400 and 500 nm decreased dramatically over blue light exposure time (Fig. 2c). The optical density (OD) at 470 nm (from Fig. 2c) versus the blue light dose can be well fitted with equation (1) (Fig. 2d). Additionally, naftifine-treated or CrtM-mutant MRSA extracts were insensitive to blue light exposure, indicated by their nearly unchanged absorption spectra (Supplementary Fig. 6a-c). These findings collectively suggest that STX is prone to bleaching under blue light irradiance.

**Fig. 2.**
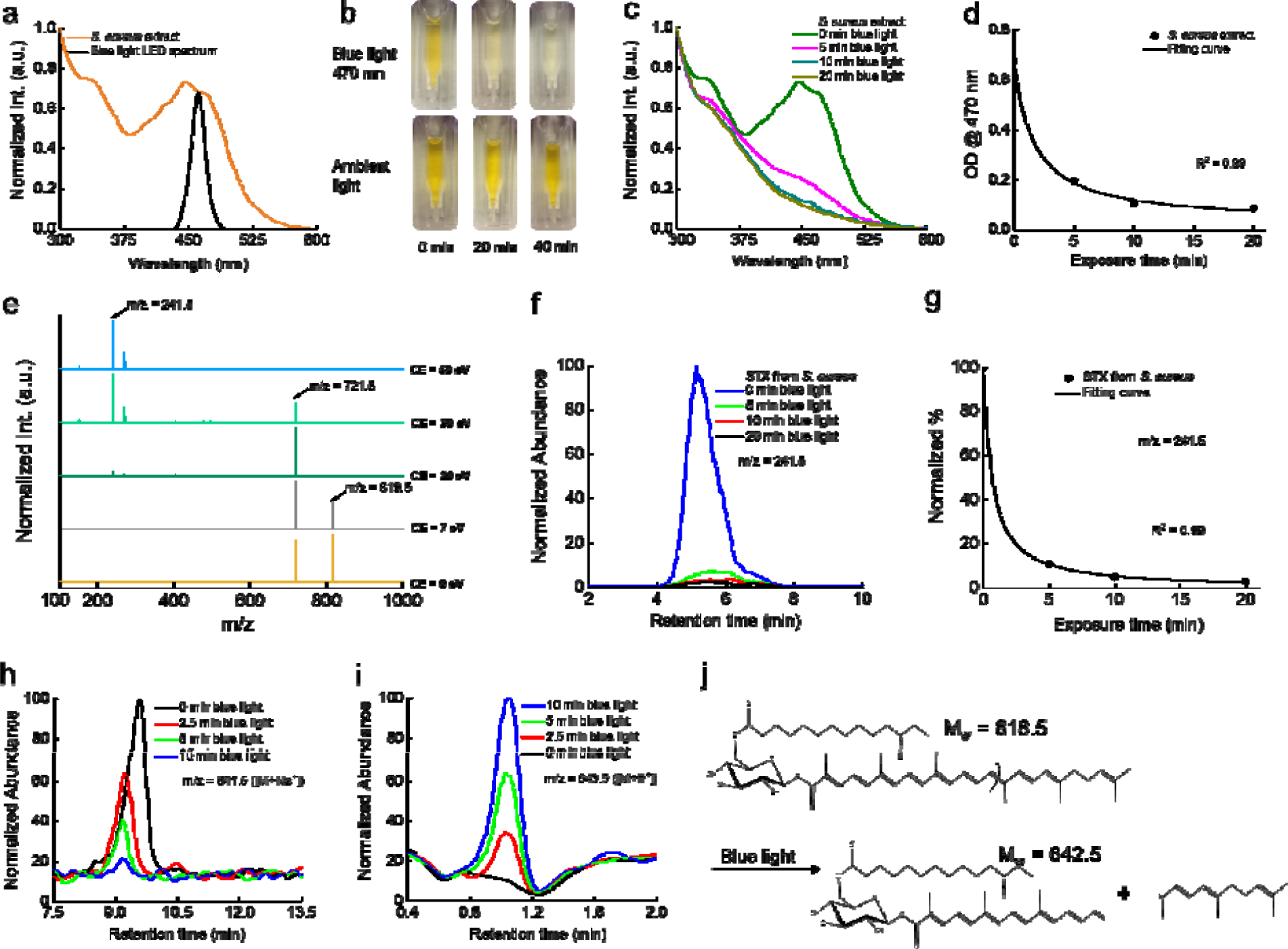
Mass spectrometry unveils photochemistry of STX under blue light exposure. (a) Absorption spectra of crude STX extract (yellow) and blue light LED (black). (b) Pictures of crude STX extract exposed to blue light and ambient light with different time intervals. (c) Absorption spectra of crude STX extract over blue light exposure time. (d) OD of crude STX extract at 470 nm adapted from (c) over blue light exposure time. Data points are fitted by equation (1). (e) MS spectra of crude STX extract at different collision energy with peaks of m/z = 819.5, m/z = 721.5 and m/z = 241.5 highlighted by black arrows. CE, collision energy. (f) HPLC spectra of STX from untreated *S. aureus* over blue light exposure time. (g) The amount of STX calculated from (f) over blue light exposure time. Quantification of STX is determined from the peak area of STX in HPLC spectra. Data points are fitted by equation (1). UPLC spectra of targeted STX (h) and its corresponding product (i) over blue light exposure. (j) Representative breakdown pathway of STX by blue light indicated by (i).

To quantitate the photobleaching process, we studied STX degradation induced by blue light irradiation by mass spectrometry (MS). Supplementary Fig. 7 presents the MS spectrum of *S. aureus* extract with m/z ranging from 200 to 1000 eV at a collision energy of 10 eV. An abundant peak appears at m/z = 721.5, while a weaker peak at m/z = 819.5 ([M+H_+_]) is consistent with the molecular weight of STX (M_w_ = 818.5 g/mol). To find out the relationship between m/z = 721.5 and m/z = 819.5, we gradually increased the collision energy from 0 to 20 eV. In Fig. 2e, the abundance of m/z = 721.5 increases relative to that of m/z = 819.5 with increasing collision energy, which indicates m/z =721.5 is a product ion from m/z = 819.5. These data also prove that STX is the major species in *S. aureus* extract. When the collision energy was higher than 30 eV, m/z = 241.5, a product of the precursor ion m/z = 721.5, became dominant and presented as a stable marker (Fig. 2e). Thus, to accurately quantify the amount of STX versus blue light dose, we targeted the peak area in high-performance liquid chromatography (HPLC) spectra specifically from ion m/z = 241.5 (Fig. 2f). Figure 2g depicts the blue light bleaching dynamics of STX. Blue light exposure for 5 min (dose: 27 J/cm^2^) decomposed 90% of STX extracted from 3.29× 10^9^ colony-forming-units (CFU/mL) *S. aureus* (Fig. 2g), and a dose of 54 J/cm^2^ bleached all the STX pigments (data not shown). In contrast, naftifine-treated and CrtM-mutant *S. aureus* extracts had negligible response to blue light exposure (Supplementary Fig. 6d-f).

Next, we employed time-of-flight MS/MS (see Methods) to elucidate how blue light decomposed STX. Different from the m/z = 819.5 peak where STX locates in the HPLC spectra, STX crests at m/z = 841.5 in the ultra-performance liquid chromatography (UPLC) spectra (Fig. 2h), which is an adjunct between STX and Na^+^. Degradation of STX would bolster the aggregation of chemical segments. Accordingly, we screened a patch of the products after STX photobleaching (Supplementary Fig. 8). In particular, the intensity of the peak at m/z = 643.5 representing an adjunct between a STX segment with H^+^, significantly increased as blue light exposure elongates (Fig. 2i). Figure 2j illustrates how this segment is formed from breakdown of one C=C bond in STX during blue light bleaching. We note that interpretation of other products (Supplementary Fig. 8a-i) necessitates further in-depth analysis.

Given STX is critical to the integrity of *S. aureus* cell membrane^19^, we questioned whether blue light alone could eradiate MRSA through bleaching STX. Blue light at 405-420 nm has been used for MRSA suppression^22^. Yet the efficacy is limited and the molecular mechanism remain elusive. Here, we show that STX is the molecular target of blue light irradiation. We find that increasing blue light dose steadily decreased the level of MRSA CFU (Fig. 3a). Moreover, MRSA was more sensitive to blue light exposure than the CrtM mutant (Supplementary Fig. 9). Nevertheless, the positive impact on the reduction of CFU did not improve when the blue light dose exceeded 216 J/cm^2^ with 60-min exposure time (Fig. 3a). To investigate the reason, we continuously monitored the growth of MRSA in fresh medium after 10-min blue light exposure. Remarkably, MRSA exposed to blue light was able to recover and multiply after being cultured in medium (Fig. 3b). Therefore, photobleaching STX alone is not sufficient to kill MRSA completely.

**Fig. 3.**
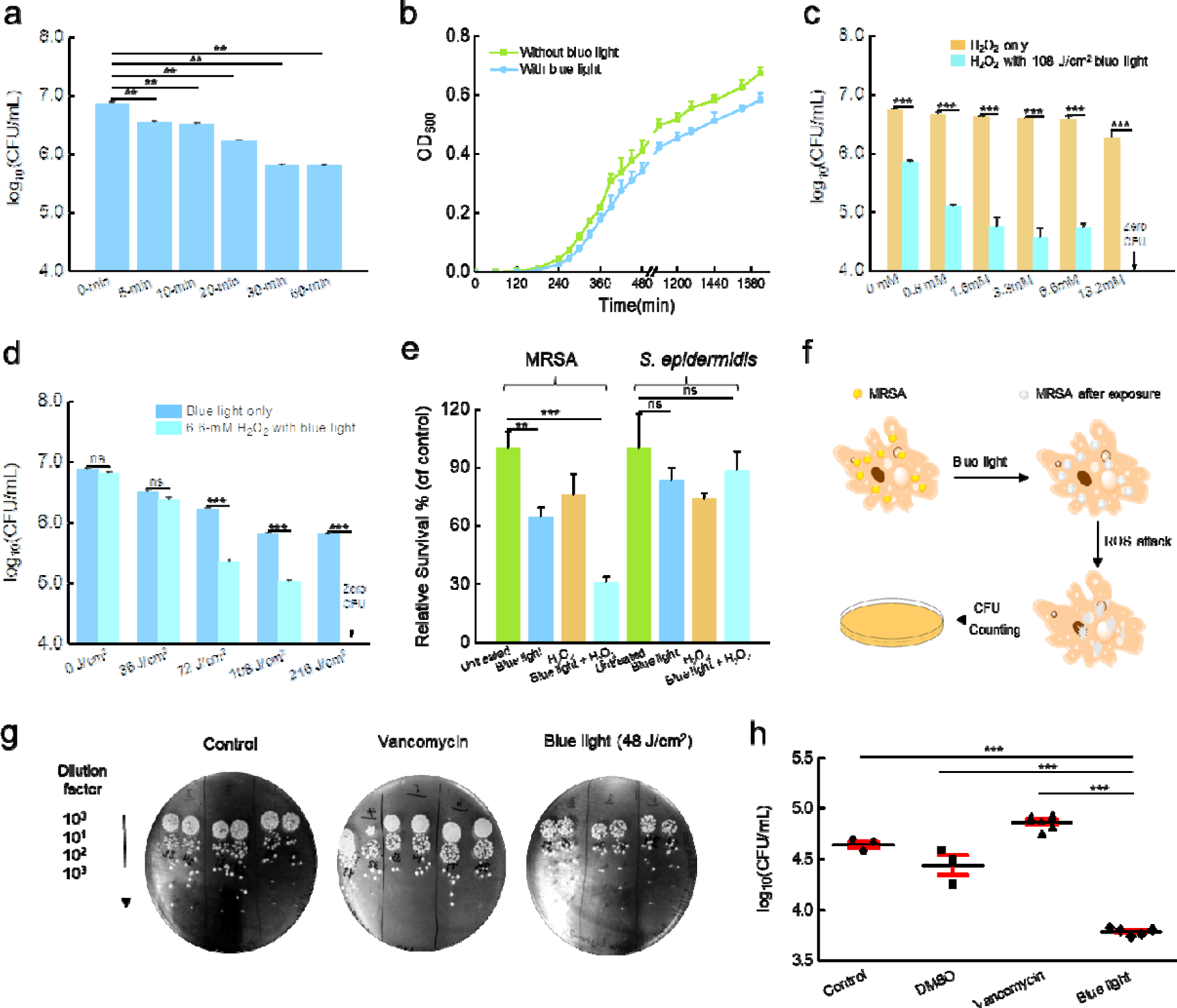
Blue light and reactive oxygen species work synergistically to eliminate MRSA in culture and in macrophages. (a) MRSA CFU under blue light exposure. Blue light intensity: 60 mW/cm. (b) Growth curves of untreated and blue light-treated MRSA (culture starts at t = 0 min immediately after the 10-min blue light exposure. Blue light dose: 120 J/cm^2^). (c) MRSA CFU of H_2_O_2_-treated and blue light plus H_2_O_2_-treated groups at different H_2_O_2_ concentrations. (d) MRSA CFU of blue light-treated and blue light plus H_2_O_2_-treated groups at different blue light doses. (e) Relative survival percentage of MRSA versus *S. epidermidis* under different treatment schemes. Blue light dose: 60 J/cm^2^. H_2_O_2_:13.2 mM, 5-min culture time. (f) Schematics illustrate the role of blue light plays in assisting macrophages to kill intracellular MRSA (not drawn to scale). (g-h) Pictures of spread plates (g) and statistical analysis (h) of CFU results of MRSA-infected macrophages from untreated, vancomycin-treated and blue light-treated groups. Error bars show standard error of the mean from at least three replicates.

Considering that STX serves as an indispensable antioxidant for MRSA, we then explored whether photobleaching of STX could sensitize MRSA to reactive oxygen species (ROS) such as hydrogen peroxide (H_2_O_2_). We examined the viability of MRSA exposed to H_2_O_2_ after blue light exposure. When MRSA was treated with blue light (dose: 108 J/cm^2^) followed by an increasing concentration of H_2_O_2_, a significant reduction (p < 0.001) in CFU was obtained (Fig. 3c). Strikingly, 108-J/cm^2^ blue light exposure combined with 13.2 mM of H_2_O_2_ (culture time: 20 min) eradicated 10^7^ MRSA CFU completely (Fig. 3c). Therefore, we hypothesized that blue light plus H_2_O_2_ work synergistically towards MRSA-killing. To verify this synergistic effect, we performed the same measurements at various blue light doses while fixing the concentration of H_2_O_2_ (Fig. 3d). Then we calculated the fractional concentration index using an established model for synergy evaluation (see Methods). A fractional concentration index of 0.45 was obtained, indicating a strong synergy between blue light and H_2_O_2_ in eradication of MRSA. Noteworthy, this treatment did not affect other species of staphylococci, such as *S. epidermidis* (Fig. 3e), that lacks carotenoids.

Studies dating back to the 1970s have demonstrated that MRSA is able to invade and survive inside mammalian cells, particularly within macrophages^23^. Though macrophages secrete small effector molecules, including ROS, bacteria including MRSA are capable of neutralizing these effector molecules by producing antioxidants such as STX^2^. Meanwhile, antibiotics are generally ineffective at clearing intracellular MRSA in large part due to the efflux of drug by phagocytic membrane, which poses an alarming threat to the host cells^23^. As we have demonstrated that blue light plus H_2_O_2_ kills MRSA synergistically, we wondered whether blue light could synergize with the ROS inside macrophage cells to eliminate intracellular MRSA as illustrated in Fig. 3f. To evaluate this point, we infected macrophage cells by incubation with MRSA for one hour. The infected macrophages were then exposed to 2-min blue light (dose: 48 J/cm^2^) twice over a 6-hour interval. The macrophages were subsequently lysed to enumerate CFU of MRSA (spread plates shown in Fig. 3g). Figure 3h compiled the statistical analysis of different groups. Notably, a nearly one-log_10_ reduction in CFU was found in the blue light-treated group in comparison to the untreated. On the contrary, vancomycin was unable to eradicate intracellular MRSA. Additionally, we found that whole blood could eradicate most of MRSA after STX bleaching by blue light (Supplementary Fig. 10). These findings collectively suggest that blue light is capable of assisting neutrophils to eradicate intracellular MRSA.

Besides residing inside host immune cells, *S. aureus* is capable of forming biofilms. Due to difficulties for antibiotics to penetrate the matrix of biofilm termed extracellular polymeric substance^24^, bacterial biofilms present a significant source of treatment failure and recurring infection in patients^24^. Compared to antibiotics, an unparalleled advantage of our photobleaching therapy lies in the fact that photons can readily penetrate through a cell membrane or a biofilm, or even a layer of tissue. To explore whether STX bleaching could eradicate *S. aureus* inside biofilm, we grew biofilms on the bottom of glass dish and then applied blue light or daptomycin (positive control) to the biofilms. Supplementary Fig. 11 shows that blue light alone (dose: 360 J/cm^2^) reduced *S. aureus* CFU by 80%. Blue light (dose: 360 J/cm^2^) plus H_2_O_2_ (13.2 mM, 20-min culture time) reduced *S. aureus* CFU by 92%. In contrast, daptomycin (5 × minimum inhibitory concentration (MIC), 24-hour culture time) only reduced *S. aureus* CFU by 70%. These results suggest an effective way to eradicate sessile bacterial cells inside biofilms.

The promising results obtained from the intracellular infection and biofilm studies led us to evaluate the efficacy of STX photobleaching in a MRSA-infected animal model. Skin infections such as diabetic foot ulceration and surgical site infections^25^ are common causes of morbidity in healthcare settings. Notably, *S. aureus* accounts for 40% of these infections^26^. To optimize the parameters for the *in vivo* experiment, we initially proved that 2-min blue light exposure (dose: 24 J/cm^2^) could cause significant reduction in survival percent of MRSA (Supplementary Fig. 12a). Then, two times antimicrobial efficiency was obtained when cultured with H_2_O_2_ (20-min culture time, 13.2 mM) subsequently. Furthermore, 5-min culture time with H2O2 after 2-min blue light exposure (24 J/cm^2^) effectively eliminated MRSA by 60% (Supplementary Fig. 12b), which facilitates us to apply treatment in MRSA-infected animal model.

To induce skin lesions in mice, we severely irritated mice skin (5 groups; 5 mice per group) by an intradermal injection containing 10^8^ CFU of MRSA USA300 (Fig. 4a), the leading source of *S. aureus* induced skin and soft tissue infections in North America^27^. Sixty hours post injection, an open wound formed at the site of infection (Fig. 4b (top)). Topical treatments were subsequently administered to each group, twice daily for three consecutive days. All the treated groups appeared healthier compared to the control group (Fig. 4b (middle)). Then, mice were humanely euthanized and wounds were aseptically removed in order to quantify the burden of MRSA in wounds (see Methods). We further examined the physiological condition of the wounds. The untreated, fusidic acid-treated (positive control), and blue light-treated groups all showed the formation of pus below the wound, most likely due to inflammatory response. In contrast, mice receiving only H_2_O_2_ or blue light plus H_2_O_2_ treatment exhibited clean wounds that were free of purulent material, swelling, and redness around the edge of the wound (Fig. 4b (bottom)). Notably, the blue light dosage applied to treat mouse wound infection was below the ANSI safety limit for skin exposure^28^.

**Fig. 4.**
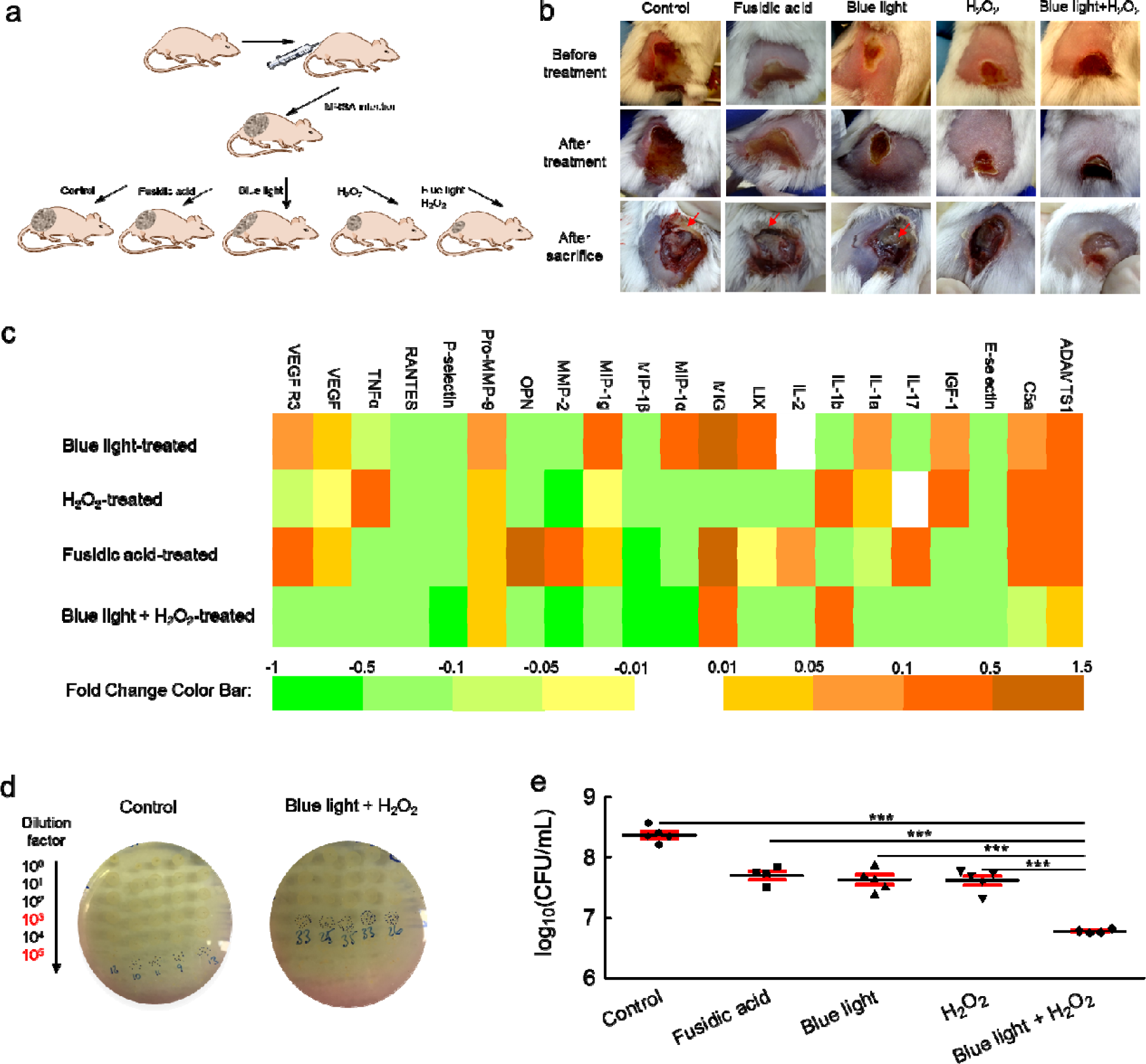
Blue light plus H_2_O_2_ effectively reduces MRSA burden in a MRSA-infected mouse wound. (a) Schematics of experiment design (not drawn to scale). (b) Pictures of mice wounds of five different groups taken before treatment, after treatment and after sacrifice. Red arrows indicate pus formation. (c) Heat map of key pro-inflammatory cytokines and markers expressed in tissue homogenate samples obtained from mice treated with blue light, H_2_O_2_, blue light plus H_2_O_2_, or fusidic acid. Orange box indicates upregulation; green box indicates downregulation; white indicates no significant change. (d) Pictures of spread plates from untreated and blue light plus H_2_O_2_-treated groups. (e) Statistical analysis of CFU results from five different groups. Error bars show the standard of the mean from N = 5 replicates. Outlier was removed through Dixon’s Q test.

To quantify the anti-inflammatory effect, we evaluated a panel of cytokines present in the supernatant of homogenized tissues extracted from the wounds of mice. By analysis of the skin homogenate collected from the MRSA mice wound model, we found the highest percent of negative fold change from around 200 kinds of cytokines in the blue light plus H2O2-treated group compared to the other groups (Supplementary Table. 1). Noteworthy, the blue light plus H2O2-treated group demonstrates the highest ratio of decreased expression of these pro-inflammatory cytokines (Fig. 4c). Specifically, there is a significant decrease observed in key pro-inflammatory cytokines (TNF-α, IL-1α, IL-2, IL-17, MIP-1α, MIP-1β, LIX) compared to the untreated mice. Furthermore, there was decreased expression of vascular endothelial growth factor receptor 3 (VEGF R3) in samples obtained from the blue light plus H_2_O_2_-treated group compared to the untreated group. This marker is overexpressed in chronic inflammatory wounds thus resulting in impaired wound reconstruction^29^. These results support a significantly decreased inflammation in the wounds of mice treated with blue light plus H_2_O_2_.

In order to quantify the burden of MRSA in wounds, homogenized wound tissue solution was inoculated onto mannitol salt agar plates (MRSA specific). Remarkably, the blue light plus H_2_O_2_-treated group showed 1.5-log_10_ reduction of CFU compared to the untreated group (Fig. 4d). Statistical analysis of CFU from the blue light plus H_2_O_2_-treated group depicted significant MRSA reduction compared to other groups (Fig. 4e). Noteworthy, the blue light plus H_2_O_2_-treated group showed one-log10 more reduction than the fusidic acid-treated group (Fig. 4e). It is worth noting that pigment is a hallmark feature of multiple pathogenic microbes^30^. Taken together, our findings show the exciting potential of treating drug-resistant bacteria by exploiting the unique photochemistry of their intrinsic pigments.

## Acknowledgements

We thank Dr. George Y. Liu of Cedars-Sinai Medical Center for providing CrtM mutant strain and Dr. Bruce Copper of Purdue University for help in the mass spectrometry study. **Funding:** This work was supported by a Keck Foundation Science & Engineering Grant to J.-X.C.

## Author contributions

P.-T.D. and X.W. made the accidental discovery. J.-X.C. conceived the concept of photobleaching therapy. J.H. and P.-T.D. mathematically analyzed the photobleaching. P.-T.D. and X.W. did the ***in vitro*** experiments. P.-T.D. and L.L. performed the biofilm experiment. H.M. and P.-T.D. carried out the intracellular study and the mice wound infection study. J.L. guided the synergy analysis. P.-T.D. and J.H. analyzed the data. M.N.S. designed the animal study and provided cytokine analysis. J.-X.C. and P.-T.D. co-wrote the paper.

## Materials and methods

### Transient absorption microscope

An optical parametric oscillator synchronously pumped by a femtosecond pulsed laser generates pump (1040 nm) and probe (780 nm) pulse trains (Supplementary Fig. 1). The pump (1040 nm) is then frequency-doubled (second harmonic generation (SHG) process) to 520 nm through a barium borate (BBO) crystal. Temporal delay between the pump and probe pulses is controlled through a motorized delay stage. The pump beam intensity is modulated with an acousto-optic modulator (AOM). The intensity of each beams is adjustable through the combination of a half-wave plate (HWP) and a polarization beam splitter (PBS). Thereafter, pump and probe beams are collinearly combined and directed into a lab-built laser-scanning microscope. Through the nonlinear process in the sample, the modulation of pump beam is transferred to the un-modulated probe beam. Computer-controlled scanning galvo mirrors are used to scan the combined laser beams in a raster scanning approach to create microscopic images. The transmitted light is collected by an oil condenser. Subsequently, the pump beam is spectrally filtered by an optical filter and the transmitted probe intensity is detected by a photodiode. A phase-sensitive lock-in amplifier then demodulates the detected signal. Therefore, pump-induced transmission changes in probe beam versus the temporal delay can be measured. This change over time delay shows different time-domain signatures of a chromophore, thus offering the origin of the chemical contrast.

### Portable blue light photobleaching apparatus

The apparatus is comprised of three parts: a blue light LED (M470L3, Thorlabs), an adjustable collimator (SM1P25-A, Thorlabs), and a power controller (LEDD1B, Thorlabs). The blue light LED has a central emission wavelength of 460 nm with a full width at half maximum of 30 nm. The beam size is adjustable through the collimator (SM1P25-A, Thorlabs). The maximal power of the blue light LED is 200 mW/cm^2^.

### Carotenoids extraction from *S. aureus* and acquisition of absorption spectrum

The pigment extraction protocol was adapted from a previous report^2^. Briefly, 100 μL of bacteria solution supplemented with 1900 μL sterile Luria-Bertani (LB) broth was cultured for 24 hours with shaking (speed of 250 rpm) at 37 °C. The suspension was subsequently centrifuged for two minutes at 7,000 rpm, washed once, and re-centrifuged. The pigment was extracted with 200 μL methanol at 55 °C for 20 minutes. Pigments from the CrtM mutant were extracted by the same method. The protocol for the treatment of *S. aureus* with naftifine was adapted from a published report^3^. Bacteria were cultured with 0.2 mM naftifine for 24 hours at 37°C with the shaking speed of 250 rpm. The extraction procedure was the same as described above. The extracted solutions were subsequently exposed to blue light (90 mW, aperture: 1 cm × 1 cm) at different time intervals (0 min, 5 min, 10 min, 20 min). Absorption spectra of the above solutions were obtained by a spectrometer (SpectraMax, M5).

### Mass spectrometry

To study the photobleaching effect on STX, we extracted crude STX from *S. aureus* and exposed the extract to blue light using the procedure described above. The separation was performed on an Agilent Rapid Res 1200 HPLC system. The HPLC-MS/MS system consisted of a quaternary pump with a vacuum degasser, thermostated column compartment, auto-sampler, data acquisition card, and triple quadrupole (QQQ) mass spectrometer (Agilent Technologies, Palo Alto, CA, USA). An Agilent (ZORBAX) SB-C8 column (particle size: 3.5 μm, length: 50 mm, and internal diameter: 4.6 mm) was used at a flow rate of 0.8 mL/min. The mobile phase A was water with 0.1% formic acid and mobile phase B was acetonitrile with 0.1% formic acid. The gradient increased linearly as follows: 5% B, from one to five min; 95% B from five to six min, and 5% B. Column re-equilibration was 6-10 min, 5% B. The relative concentration of STX was quantified using MS/MS utilizing the Agilent 6460 QQQ mass spectrometer with positive electrospray ionization. Quantitation was based on multiple reaction monitoring. Mass spectra were acquired simultaneously using electrospray ionization in the positive modes over the m/z range of 100 to 1000. Nitrogen was used as the drying flow gas.

In order to understand how STX degrades when exposed to blue light, an Agilent 6545 quadrupole time-of-flight (Q-TOF) (Agilent, Santa Clara, CA, USA) was exploited to conduct the separation and quantification steps. This ultra-performance liquid chromatography (UPLC)-MS/MS utilized an Agilent (ZORBAX) SB-C8 column (particle size: 3.5 μm, length: 50 mm, and internal diameter: 4.6 mm) to conduct the separation at a flow rate of 0.8 mL/min. The relative concentration of STX was quantified using MS/MS utilizing the Agilent 6545 Q-TOF MS/MS with positive electrospray ionization. The mobile phase was composed of water (A) and acetonitrile (B). The gradient solution with a flow rate of 0.8 mL/min was performed as follows: 85% B, from 0 to 30 min; 95% B, from 30 to 31 min; 85% B, from 31 to 35 min; 85% B, after 35 min. The sample injection volume was 20 μL The UPLC-MS/MS analysis was performed in positive ion modes in the m/z range of 100-1100.

### *In vitro* assessment of synergy between blue light and H_2_O_2_

MRSA USA300 was cultured in sterile LB broth in a 37°C incubator with shaking (at 250 rpm) until the suspension reached the logarithmic growth phase (OD_600_ = 0.6). Thereafter, an aliquot (20 μL) of the bacterial suspension was transferred onto a glass slide. Samples were exposed to blue light at different time lengths and variable light intensities. For groups treated with H_2_O_2_, bacteria were collected in either LB or phosphate buffered saline (PBS) supplemented with H2O2 at different concentrations (0 mM, 0.8 mM, 1.6 mM, 3.3 mM, 6.6 mM, and 13.2 mM). The solutions were cultured for 20 min. The solution was serially diluted in sterile PBS and transferred to LB plates in order to enumerate the viable number of MRSA CFU. Plates were incubated at 37 °C for 24 hours before counting viable CFU/mL. Data are presented as viable MRSA CFU/mL and percent survival of MRSA CFU/mL in the treated groups.

### Fluorescence mapping of live and dead *S. aureus* in biofilm

An overnight culture of *S. aureus* (ATCC 6538) was grown in a 37 °C incubator with shaking (at 250 rpm). Poly-D-lysine (Sigma Aldrich) was applied to coat the surface of glass bottom dishes (35 mm, In Vitro Scientific) overnight. The overnight culture of *S. aureus* was diluted (1:100) in LB containing 5% glucose and transferred to the glass bottom dishes. The plates were incubated at 37°C for 24-48 hours in order to form mature biofilm. Thereafter, the media was removed and the surface of the dish was washed gently with sterile water to remove planktonic bacteria. Plates were subsequently treated with blue light alone (200 mW/cm^2^, 30 min), H_2_O_2_ (13.2 mM, 20 minutes) alone, or a combination of blue light and H_2_O_2_. Groups receiving H_2_O_2_ were quenched through addition of 0.5 mg/mL catalase (Sigma Aldrich, 50 mM, pH = 7 in potassium buffered solution). After treatment, biofilms were immediately stained with fluorescence dyes, as follows.

To confirm the existence of biofilm on the glass bottom surface, a biofilm matrix stain (SYPRO^®^ Ruby Biofilm Matrix Stain, Invitrogen) was utilized. Biofilms were stained with the live/dead biofilm viability kit (Invitrogen) for 30 minutes to quantify the survival percent of *S. aureus* in the biofilm after treatment. The biofilms were washed with sterile water twice and then imaged using a fluorescence microscope (OLYMPUS BX51, objective: 60x, oil immersion, NA = 1.5). Two different excitation channels (live: FITC; dead: Texas Red) were utilized in order to map the ratio of live versus dead cells within the biofilm. The acquired images were analyzed by ImageJ. Statistical analysis was conducted via a two-paired t-test through GraphPad Prism 6.0 (GraphPad Software, La Jolla, CA).

### Intracellular MRSA infection model

Murine macrophage cells (J774) were cultured in Dulbecco’s Modified Eagle Medium (DMEM) supplemented with 10% fetal bovine serum (FBS) at 37 °C with CO_2_ (5%). Cells were exposed to MRSA USA400 at a multiplicity of infection of approximately 100:1. 1-hour post-infection, J774 cells were washed with gentamicin (50 μg/mL, for one hour) to kill extracellular MRSA. Vancomycin, at a concentration equal to 2 μg/mL (4 × MIC, MIC: minimal inhibition concentration), was added to six wells. Six wells received blue light treatment twice (six hours between treatments) for two minutes prior to addition of DMEM + 10% FBS. Three wells were left untreated (medium + FBS) and three wells received dimethyl sulfoxide at a volume equal to vancomycin-treated wells. Twelve hours after the second blue light treatment, the test agents were removed; J774 cells were washed with gentamicin (50 μg/mL) and subsequently lysed using 0.1% Triton-X 100. The solution was serially diluted in PBS and transferred to tryptic soy agar plates in order to enumerate the MRSA CFU present inside infected J774 cells. Plates were incubated at 37 °C for 22 hours before counting viable CFU/mL. Data are presented as log_10_(MRSA CFU/mL) in infected J774 cells in relation to the untreated control. The data was analyzed via a two-paired *t*-test, utilizing GraphPad Prism 6.0 (GraphPad Software, La Jolla, CA).

### *In vivo* MRSA mice wound model

All animal experiments were conducted following protocols approved by Purdue Animal Care and Use Committee (PACUC). To initiate the formation of a skin wound, five groups (N = 5) of eight-week-old female Balb/c mice (obtained from Harlan Laboratories, Indianapolis, IN, USA) were disinfected with ethanol (70%) and shaved on the middle of the back (approximately a one-inch by one-inch square region around the injection site) one day prior to infection as described from a reported procedure (Ref. 31). To prepare the bacterial inoculum, an aliquot of overnight culture of MRSA USA300 was transferred to fresh tryptic soy broth and shaken at 37 °C until an OD_600_ value of ~1.0 was achieved. The cells were centrifuged, washed once with PBS, re-centrifuged, and then re-suspended in PBS. Mice subsequently received an intradermal injection (40 μL) containing 2.40 × 10^9^ CFU/mL MRSA USA300. An open wound formed at the site of injection for each mouse, ~60 hours post-infection.

Topical treatment was initiated subsequently with each group of mice receiving the following: fusidic acid (2%, using petroleum jelly as the vehicle), 13.2 mM H_2_O_2_ (0.045%, two-minute exposure), blue light (two-minute exposure, 24 J/cm^2^), or a combination of blue light (two-minute exposure) and 13.2 mM H_2_O_2_ (two-minute exposure). One group of mice was left untreated (negative control). Each group of mice receiving a particular treatment regimen was housed separately in a ventilated cage with appropriate bedding, food, and water. Mice were checked twice daily during infection and treatment to ensure no adverse reactions were observed. Mice were treated twice daily (once every 12 hours) for three days, before they were humanely euthanized via CO_2_ asphyxiation 12 hours after the last dose was administered. The region around the skin wound was lightly swabbed with ethanol (70%) and excised. The tissue was subsequently homogenized in PBS. The homogenized tissue was then serially diluted in PBS before plating onto mannitol salt agar plates. Plates were incubated for at least 19 hours at 37 °C before viable MRSA CFU/mL were counted for each group. Outlier was removed based upon the Dixon Q Test. Data were analyzed via a two-paired t-test, utilizing GraphPad Prism 6.0 (GraphPad Software, La Jolla, CA).

### Statistical analysis

Data were present as mean values and its standard error of the mean. Statistical analysis was conducted through two-paired *t*-test. *** means significantly different with the *p*-value < 0.001. ** means significantly different with the *p*-value < 0.01. * means significantly different with the *p*-value < 0.05. ns means no significance.

### Human whole blood

After photobleaching of MRSA by blue light, MRSA were either cultured in sterile PBS (control) or human whole blood (Innovative Research Inc., Novi, MI) for 9 hours. The efficacy was evaluated through enumerating CFU.

### Photobleaching model

To analyze the time-lapse transient absorption signals, we utilized a mathematical model which was originally used to depict the photobleaching of photosensitizers happening during a photodynamic process^16^:

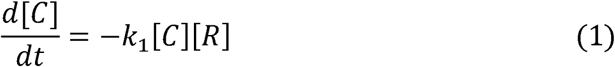

where *t* is the duration time, [C] is the concentration of chromophore (e.g., carotenoids in *S. aureus*),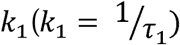 is the rate constant of first-order photobleaching with *τ*_1_ being the first-order photobleaching time constant, [R] is the concentration of active agents (the chromophores which have interaction with light):

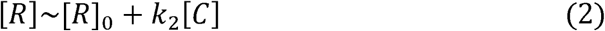

where 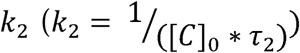 is the rate constant of second-order photobleaching with *τ*_2_ being the second-order photobleaching time constant, [*R*]_0_ is the initial concentration of the active agent, respectively. The combination of equation (1) and equation (2) leads to:

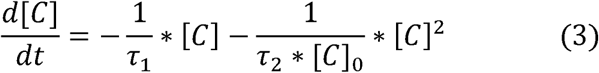

The solution for equation (3) is:

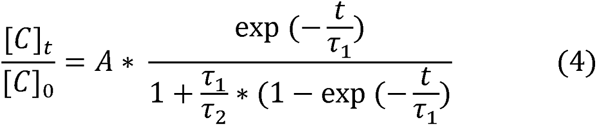

where A is a constant. When first-order photobleaching process pivots (usually happening for low concentration of chromophore and having the involvement of oxygen), *τ*_2_ → ∞, then equation (4) becomes:

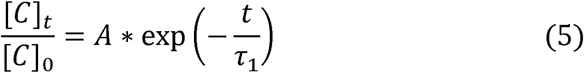

which is similar to first-order kinetic reaction. At this occasion, the photobleaching rate is linearly proportional to the concentration of chromophore. When second-order photobleaching process dominates (usually happening for high concentration of chromophore potentially through triplet-triplet annihilation), *τ*_1_ → ∞, then equation (4) becomes:

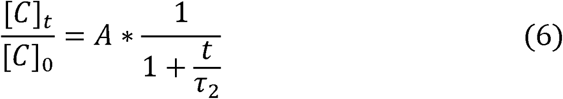

Under this condition, the photobleaching rate is proportional to the square of chromophore concentration. According to the fitting result, *S. aureus* belongs to second-order bleaching with *τ*_1_ → ∞.

### Synergistic antimicrobial effect analysis

The synergy analysis is based on the calculation of fractional bactericidal concentration index (FBCI) (Ref. 32). Here, FBC stands for fractional bactericidal concentration and MBC is minimal bactericidal concentration. FBCI was calculated as follows: FBC of drug A = MBC of drug A in combination with drug B/MBC of drug A alone, FBC of drug B = MBC of drug B in combination with drug A/MBC of drug B alone, and FBCI index = FBC of drug A+FBC of drug B. An FBCI of ≤ 0.5 is considered to demonstrate synergy. Additive was defined as an FBCI of 1.

Antagonism was defined as an FBCI > 4. Since 200 mW/cm^2^ blue light did not kill 100% MRSA after 1-hour exposure time, we have

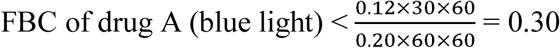

Since we found that 88 mM H_2_O_2_ is needed to eradicate all the bacteria, we have

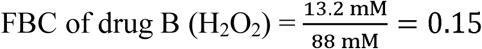

Therefore, FBCI = FBC of blue light + FBC of H_2_O_2_ < (0.30 + 0.15 = 0.45).

